# Genomic Insights into the Ecologies of Atlantic Oysters *Crassostrea virginica*

**DOI:** 10.64898/2026.10.13.758928

**Authors:** Liam Cawood, Shiying Wang, Eli Kleiman, Zhiyuan Chen, Isabel Kleiman, Nicole Fair, Ruby Brownell, Irina Tikhonova, Zicheng Li, Josephine Hoh

## Abstract

Oysters have been increasingly used as model organisms to understand how living beings interact with their environments. The eastern oyster, *Crassostrea virginica*, is a foundation species in coastal ecosystems along Northeast America, and is also rich in nutrients for human consumption. However, the labor at the oysters’ farms is intensive and time-consuming, and yields are often unpredictable and random, resulting in over-farming and disrupting the ecosystem. To overcome the unpredictability, we propose a potential farming strategies that combines current practices with genetic information derived from modern genomic technologies and high-power computational methods. This study aims to ascertain new genetic information of *C. virginica* through harvesting oysters and constructing DNA genomes. Sequence analysis provides information of genetic variation among *C. virginica*. Among several notable findings, a region on chromosome 1 harbors a long-range expansion of the innate immunity genes, which potentially illuminates a survival mechanism for *C. virginica* under constant changes of the environment. Gene expressions of the human homologs also have been reported to be significantly upregulated in immunocompromised patients who undergone the 2′3′-cUA treatment. This study offers a novel platform for eco-friendly and efficient oyster breeding and farming strategies.

## Introduction

Marine invertebrates thrive in highly dynamic coastal environments, yet the genomic basis of their resilience remains poorly understood. The eastern oyster (*Crassostrea virginica*) is native to the Atlantic coast of North America and plays both major economic and ecological roles. Oyster reefs stabilize shorelines, provide habitat for diverse species, and improve water quality through filter-feeding^1–3^. Because of these functions, *C. virginica* has become a focal species in restoration and aquaculture, and a model for studying the genetic basis of adaptation to environmental stress^4–7^. As a sessile filter feeder, *C. virginica* endures large fluctuations in salinity^8,9^, temperature^10,11^, oxygen^12,13^, and pathogen load^6,7,14^. These persistent stressors are thought to drive lineage-specific expansions of immune and stress-response gene families^15,16^. Selective breeding and genomic resources have reinforced its importance as a research model^17–19^, yet fully resolving the mechanisms of gene family diversification requires an accurate reference genome.

The current assembly (*C_virginica_3.0*) contains haplotig redundancy and chromosomal misassemblies^20^, limiting analyses of gene family diversification and distorting cross-species comparisons^16^. With growing interest in comparative genomics across oysters, such as *Magallana gigas*, *M. hongkongensis*, and *Ostrea edulis*^21–23^, a structurally accurate *C. virginica* genome is essential for reliable evolutionary inferences. This species occupies a pivotal position within the *Crassostreinae* subfamily, having diverged from Pacific oysters and other bivalves over 80 million years ago^24^. This deep evolutionary split, combined with the species’ remarkable resilience to fluctuating estuarine environments, makes *C. virginica* a compelling model for investigating how molluscan genomes evolve under diverse ecological pressures.

In this study, we present a high-quality chromosome-level assembly that corrects previous errors and reveals new evolutionary insights. Notably, we show that genome divergence among oysters is dominated by extensive intrachromosomal rearrangements. We also identified a 13Mb lineage-specific expansion enriched for immune and stress-response families in *C.virginica*. Together, these findings show that structural variation and lineage-specific expansion interact to generate unusually large immune clusters, providing new insight into how genome remodeling shapes adaptation in marine invertebrates.

## Results

### Summary

The newly assembled genome achieves 99.7% genome completeness and reduces duplicated single copy ortholog content from 18.7% to 1.1%, resolving chromosomal misassembles and eliminating haplotigs. Conservatively around 35k SNPs were found in 19 oysters vs. the reference. Comparative alignment with the genome of *Magallana gigas* (Pacific Oyster) reveals structural variation arising from intra-chromosomal rearrangements such as inversions and localized block shuffling. Most notably, we identified a 13Mb lineage-specific expansion on chromosome 1-a large, contiguous region. This 13 Mb segment harbors an immune gene family, such as ferric chelate reductase 1 (FRRS1). Phylogenetic analysis indicates modular reshaping through domain loss, retention, and diversification.

### A structurally resolved chromosome-level assembly for *C. virginica*

We assembled a high-quality chromosome-level genome of C. virginica using PacBio HiFi, Illumina WGS, and Hi-C data (Figure 1A). The final assembly contains 10 chromosomes totaling 529.0 Mb, composed of 81 contigs and 71 gaps distributed across chromosomes, with a scaffold N50 of 55.3 Mb and a contig N50 of 16.0 Mb, respectively. Analysis of 21-mers confirmed high consensus accuracy of the assembly (Figure 1B). Compared with the previous reference (*C_virginica_3.0*), our assembly is shorter but substantially more contiguous with reduced genome redundancy (BUSCO duplication from 18.7% to 1.1%) while achieving 99.7% gene completeness (Figure 1C). Hi-C contact maps confirmed chromosome-scale integrity, showing strong intra-chromosomal signals and no evidence of inter-chromosomal misjoins (Figure 1D). Whole-genome alignments indicated broad sequence consistency (80.6%) at the kilobase scale (≥1kb alignment), but divergence (40.8%) at longer alignment scales (≥10kb alignment) pointed to structural discrepancies rather than sequence-level errors (Figure 1E).

**Figure 1.**
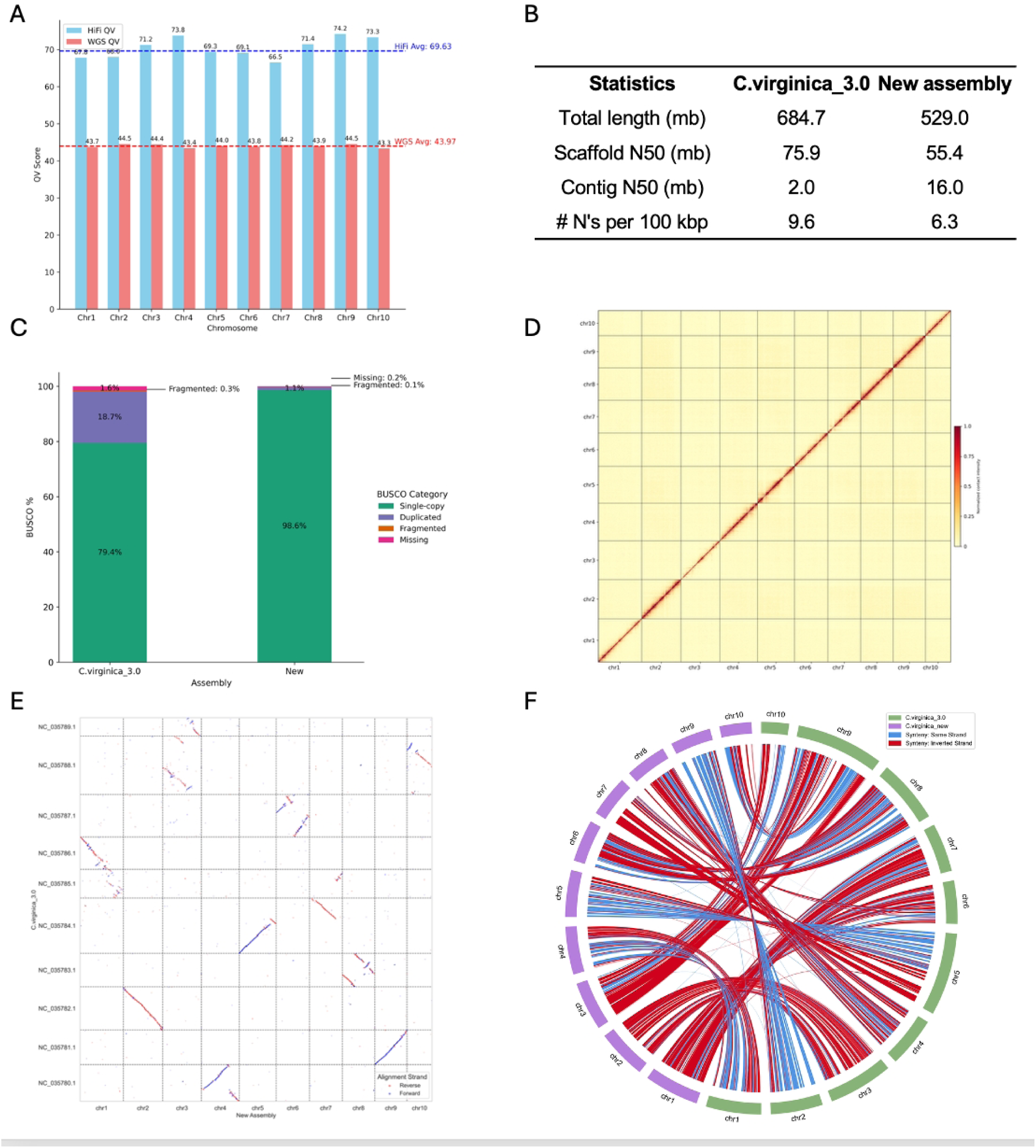
Comparison of the newly assembled C. virginica genome to the existing reference genome. **(A)** Per-chromosome 21-mer QV scores from both HiFi and Illumina WGS reads. **(B)** Summary statistics of the new assembly and *C. virginica* v3.0. **(C)** BUSCO gene completeness assessment showing increased single-copy completeness in the new assembly. **(D)** Hi-C contact map of a representative scaffold confirms chromosomal structure and accurate contig joins. **(E)** Dot plot alignment between the new and previous genomes reveals large-scale misassemblies corrected in this version. **(F)** Genome-wide synteny chordgram showing chromosomal correspondence and structural rearrangements between assemblies.

Finally, synteny alignment confirmed that several major misassemblies in cv3.0, including the fragmentation of chromosome 1, fusion of chromosomes 5/7, and misjoins involving chromosomes 3, 5, and 10, are corrected in our assembly (Figure 1F).

### Genome architecture and functional annotation

We annotated the *C. virginica* genome to characterize sequence composition and repetitive content (Figure 2A). Illumina WGS coverage was uniform (mean 30.8×, SD ±7.74×), with only 0.04% of windows deviating more than ±2 standard deviations, indicating consistent representation across the genome. Repeat annotation using a custom library identified 43.2% of the genome as repetitive, including DNA transposons (7.1%), LTRs (7.1%), LINEs (3.4%), and SINEs (3.6%) (Table S1). Telomeric repeats were detected near the ends of six chromosomes, consistent with partial telomeric structures (Table S2). Putative centromeric arrays were identified on seven chromosomes, spanning 10-80kb with high sequence similarity among repeat units (Table S3). Their positions coincide with TE-rich regions and match prior cytogenetic centromere estimates^25^ (Table S4), supporting centromeric identity. Gene prediction identified 33,259 protein-coding genes (Figure 2B, Data S1), of which around 68% received functional annotation (Figure 2C). Gene density negatively correlated with transposable element content (Figure 2D-E), consistent with repeat accumulation in gene-poor regions. Together with analyses of recent segmental duplications (Figure 2F), this annotation provides a robust framework for comparative and functional studies of genome structure and evolution.

**Figure 2.**
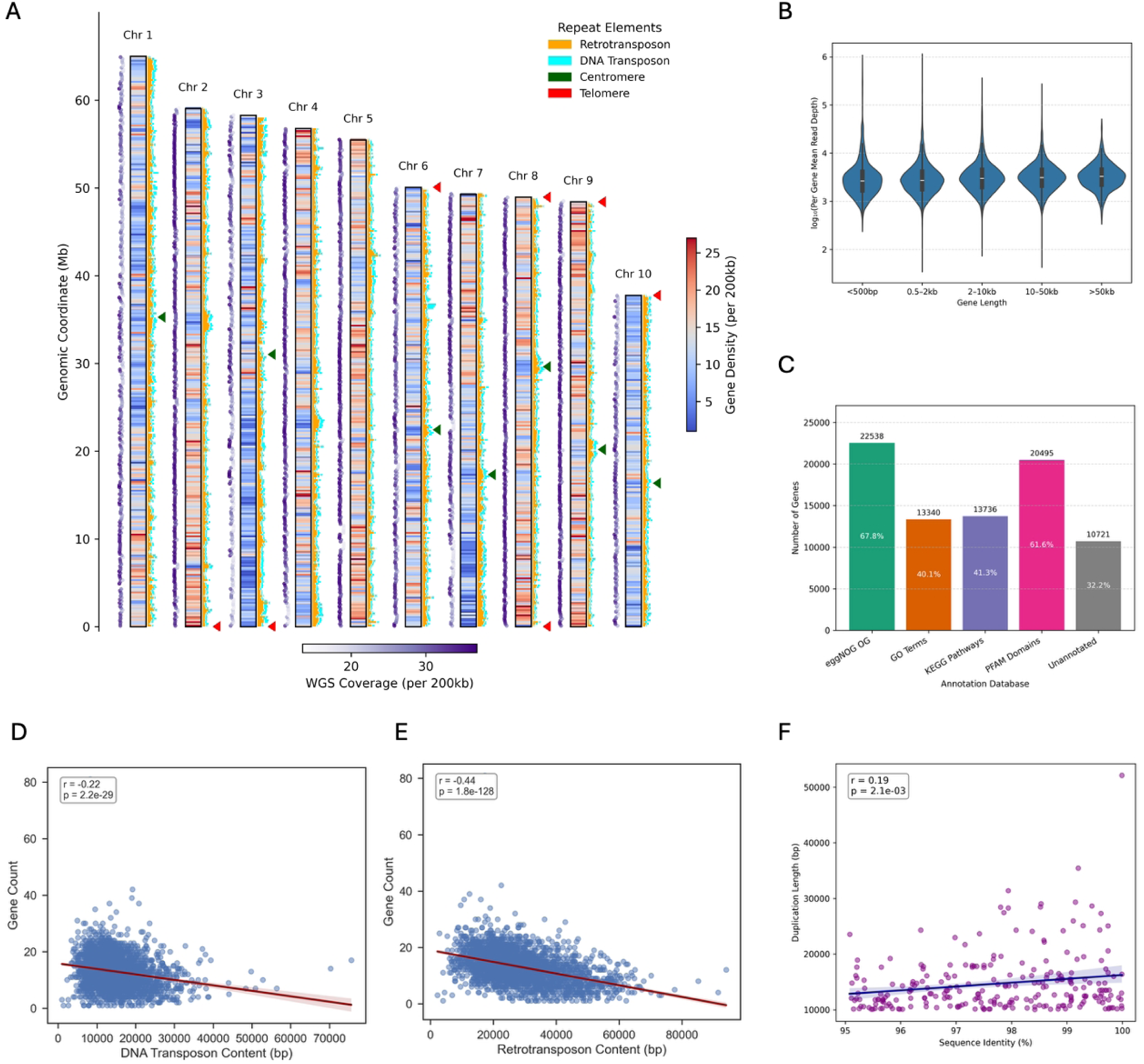
Genome-wide landscape and annotation evaluation for the *Crassostrea virginica* reference genome. **(A)** Chromosomal landscape of repeat content, gene density, and sequencing coverage across the 10 chromosomes of *Crassostrea virginica*. Each vertical bar represents a chromosome scaled by length, with the internal heatmap indicating gene density per 200 kb window. To the left of each chromosome, WGS coverage is plotted as a purple gradient. To the right, transposable elements per 200kb window are shown as stacked bars while putative centromere and telomere arrays are indicated by triangles. **(B)** Violin plot showing gene model support (mean RNA-seq read depth per gene) across gene length bins. **(C)** Functional annotation statistics for predicted protein-coding genes. Colored bars indicate genes with eggNOG orthogroup, GO term, KEGG pathway, and PFAM domain annotations; the unannotated portion is shown in gray. **(D–E)** Scatterplots illustrating negative correlations between gene count and TE content per 200 kb window **(F)** Scatterplot showing a weak but significant positive correlation between duplication length and sequence identity among recent large segmental duplications (≥10 kb, ≥95% identity). Each dot represents a duplication, colored in purple with a regression line showing the correlation.

### Genome-wide SNP variation across individuals

To provide a preliminary view of genome-wide polymorphisms, we analyzed WGS data from 19 additional *C. virginica* samples. After stringent filtering, we identified 34,810 high-confidence SNPs defined as loci where all 19 individuals differed from the reference allele. These variants were distributed across all 10 chromosomes, with counts ranging from 570 on chromosome 4 to 5,546 on chromosome 1 (Table S5). As expected, all non-reference samples exhibited exclusive alternate homozygosity at these loci, with per-sample totals ranging from 30,530 to 32,558 (Table S6). While the modest sample size and conservative SNP definition likely underestimate true genome-wide variation, this analysis demonstrates the utility of the new reference for population-level variant discovery.

### Chromosome stability and evolutionary divergence across oysters

To place the *C. virginica* genome in an evolutionary context, we compared it with three chromosome-level oyster assemblies with available gene annotations: *O. edulis*, *M. gigas*, and *M. angulata*. Whole-genome synteny revealed strict one-to-one correspondence across all ten chromosomes (Figure 3A), but synteny block size differed: *O. edulis* showed significantly longer blocks than *C. virginica*, whereas block size distributions were similar between *C. virginica* and *Magallana* species (Figure 3B). Chromosomes 3 and 10 exhibited markedly lower conservation (Figure 3C), coinciding with their high repeat content (Table S1), suggesting repeat accumulation may underlie structural divergence. Orthogroup analysis recovered 14,719 gene families shared by all four species, with *C. virginica* harboring the largest number of species-specific orthogroups (n=1,049) (Figure 3D). Phylogenetic analysis revealed a long branch leading to *C. virginica* (Figure 3E), indicating an accelerated amino acid substitution rate. *C. virginica* exhibited the largest number of expanded gene families among the four species (Figure 3F), further indicating its lineage-specific diversification.

**Figure 3.**
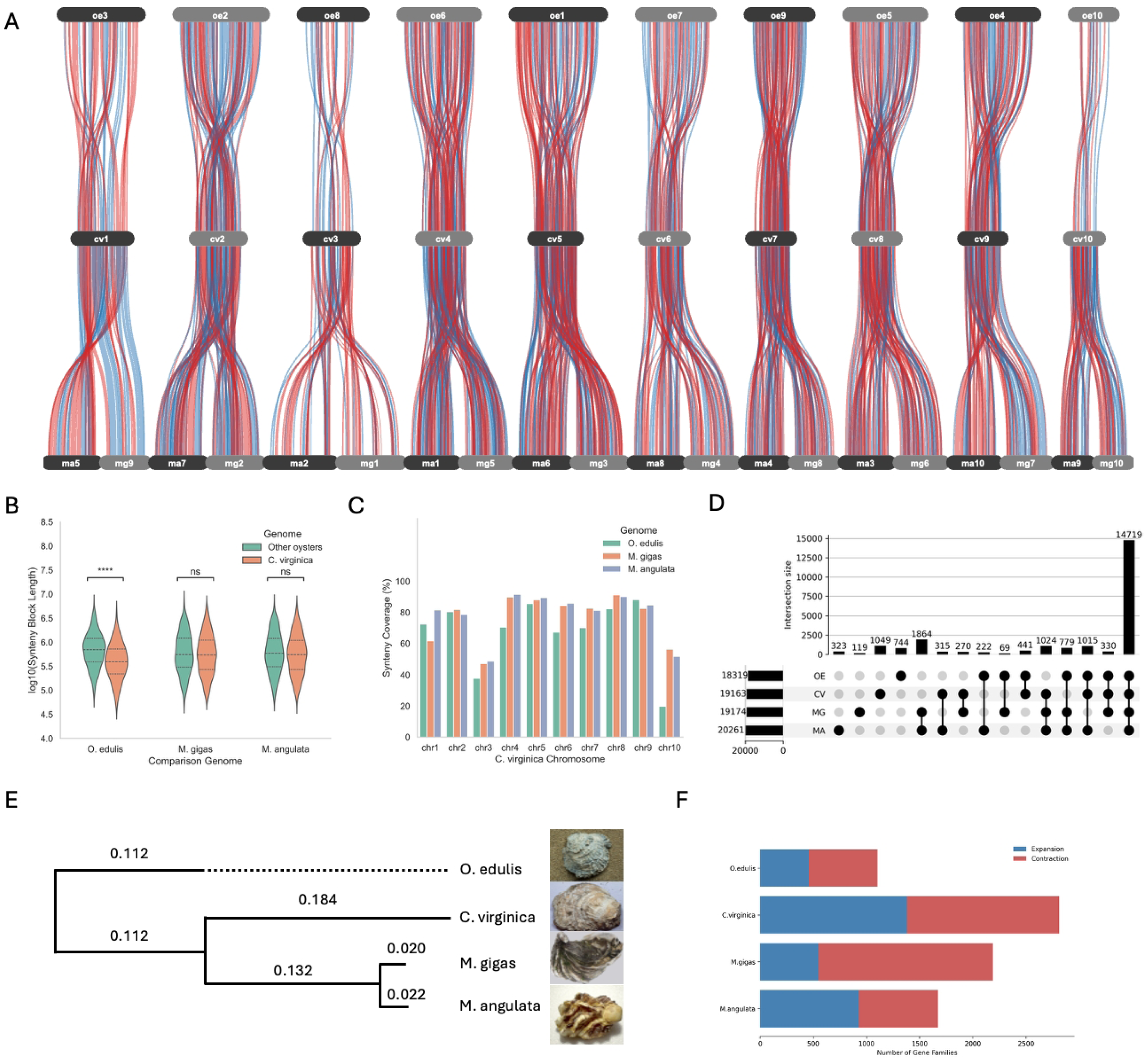
Comparative genome structure and orthogroup evolution among four chromsome level oyster genomes. **(A)**Synteny alignment between *C. virginica* and three other oyster genomes. Each ribbon represents a conserved synteny block between chromosomes, with blue and red indicating forward and reverse orientation, respectively. **(B)**Violin plots comparing synteny block lengths (log10-transformed) between *C. virginica* and each comparison genome. **(C)**Bar plot showing the percentage of each *C. virginica* chromosome covered by syntenic blocks with each comparison genome. *C. virginica* chromosomes 3 and 10 consistently exhibit reduced synteny coverage across species. **(D)**UpSet plot showing the distribution of orthogroups shared among the four species. **(E)**Species tree based on orthologous proteins, with branch lengths representing relative amino acid substitution rates. **(F)**Stacked bar plot of the number of significantly expanded (blue) and contracted (red) gene families inferred by CAFE5 in each species.

### Fine-scale chromosomal remodeling shapes divergence between *C. virginica* and *M. gigas*

To further investigate the genomic basis of divergence in *C. virginica*, we performed a detailed pairwise comparison with *M. gigas*, a well-studied Pacific oyster species. As an ecologically distinct yet phylogenetically close relative, *M. gigas* provides an ideal evolutionary contrast for assessing genome-scale conservation, structural rearrangement, and gene content variation within the *Crassostrea* lineage. After filtering for non-overlapping blocks with ≥10 colinear genes, we identified 348 synteny blocks spanning 69.6% of the *C. virginica* genome and 61.3% of the *M. gigas* genome (Data S2), comprising 11,394 orthologous gene pairs. Genome-wide visualization revealed extensive chromosome-level collinearity, with no inter-chromosomal translocations detected (Figure 4A).

**Figure 4.**
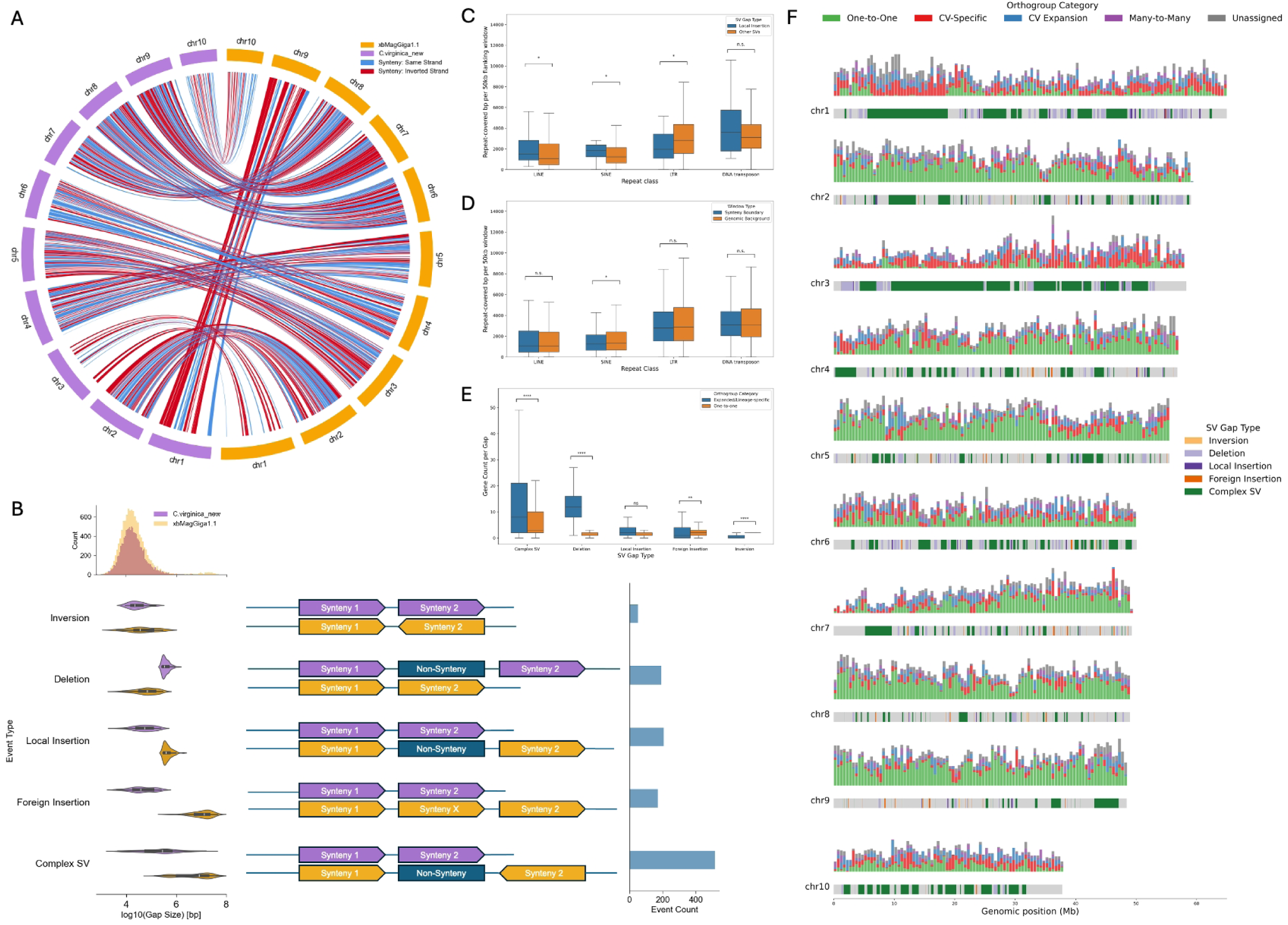
Comparative synteny and structural variation between *C. virginica* and *M. gigas*. **(A)**Genome-wide synteny alignment between *C. virginica* and *M. gigas*, visualized as inter-chromosomal chords. Blue chords represent forward-strand (collinear) alignments, while red chords indicate reverse-strand (inverted) syntenic blocks. **(B)**Structural variation classification based on synteny breakpoints. Top: genome-wide distribution of gap sizes between adjacent anchor gene pairs. Left: violin plots showing the gap length distribution associated with each structural variation type. Right: bar plot summarizing the number of detected events per SV class. **(C)**Transposable element content in 50 kb windows flanking synteny boundaries, grouped by repeat class. Total repeat-covered base pairs are shown for each TE class. **(D)**TE enrichment near synteny breakpoints compared to genome-wide background levels. Bars represent average repeat coverage in boundary-proximal windows versus randomly sampled genome-wide regions. **(E)**Gene orthogroup content within local SV gaps (insertions, deletions, and inversions). The distribution of one-to-one, expanded, and species-specific orthogroups is shown per event, highlighting differential enrichment patterns by SV type. **(F)**Chromosome ideograms summarizing orthogroup distribution and SV gap types. Each stacked bar represents a 500 kb window across the chromosome. The top track shows the count and category of genes per window. The bottom track depicts SV-associated gaps, color-coded by event type. Gray segments indicate terminal gaps (adjacent to chromosome ends) and conserved syntenic regions between anchor pairs.

Despite conserved chromosome-scale synteny, *C. virginica* and *M. gigas* differ markedly in intrachromosomal organization (Table S7). Chromosomes 1 and 3 harbor large inversions spanning >20 Mb, with strongly reversed syntenic order (*Corr_Spearman_*=-0.95 and-0.67, respectively). Chromosomes 2, 4, 5, 6, and 8 contain numerous orientation flips (≥17), reflecting fragmented synteny and frequent local rearrangements. In contrast, chromosomes 7, 9, and 10 show very low gene order correlation (|*Corr_Spearman_* |<0.35), consistent with complex, mosaic rearrangements. These patterns indicate that intrachromosomal inversions of varying scale have been a major driver of divergence between the two species, which was estimated to split around 83 Mya^24^.

We further characterized structural variants (SVs) using an anchor gene-based approach (see Methods) to examine structural divergence beyond collinearity (Data S3). Using *C. virginica* as the anchor genome, complex SVs were most frequent (n=516), followed by local insertions (n=206), foreign insertions (n=172), deletions (n=192), and inversions (n=51). Foreign insertions were significantly larger than local insertions (*p*-value=1.8×10^-31^) (Figure 4B), consistent with their embedding of syntenic blocks. These results indicate that large insertions and complex rearrangements are the dominant contributors to collinearity disruption between *C. virginica* and *M. gigas*. Repeat content in 50 kb flanking regions of synteny breakpoints was profiled to investigate whether transposable elements contribute to genome rearrangements (Data S4). Breakpoints of local-insertion-type SVs were significantly enriched for LINEs (*p*-value=0.026), LTRs (*p*-value=0.016), and to a lesser extent SINEs (*p*-value=0.026) (Figure 4C). In contrast, breakpoints across all SVs showed repeat composition similar to the genomic background, with only modest SINE depletion (*p*-value=0.016) (Figure 4D). These results suggest that TE activity may contribute in a variant-type-specific manner to SV formation, shaping oyster genome architecture.

We heuristically classified genes into orthogroups across *C. virginica* and *M. gigas* (Data S5) to investigate ortholog discrepancies between these two species. We identified 13,183 strict one-to-one orthologs, along with 1,479 and 834 lineage-specific expansions in *C. virginica* and *M. gigas*, respectively. Additionally, 1,463 CV-specific and 825 MG-specific genes were detected, plus 690 genes in complex many-to-many groups. Within SV-associated gaps, expanded and lineage-specific genes were significantly overrepresented (Figure 4E). In particular, complex SVs and deletion gaps were enriched for non-conserved orthogroups relative to one-to-one orthologs (*p*-value=9.93×10^-5^ and 2.12×10^-27^). These patterns suggest that SVs are disproportionately associated with dynamic gene families, highlighting their differential roles in shaping genome plasticity. To provide a genome-wide view of how SVs and gene family evolution intersect across chromosomes, we overlaid orthogroup categories with SV-associated non-syntenic gaps in 500 kb windows to examine how SVs and gene family evolution intersect across chromosomes (Figure 4F). The ideogram reveals pronounced chromosomal heterogeneity in the distribution of conserved orthologs. Chromosomes 1 and 3 show the lowest mean proportions of one-to-one orthologs (0.17 and 0.13, respectively) and the highest coefficients of variation (0.97 and 1.27, respectively), coinciding with regions of major structural rearrangements (Table S8). These patterns suggest that large-scale SVs may erode conserved gene order and promote lineage-specific family turnover. In contrast, chromosomes 8 and 9 display high and stable conservation (mean proportions of one-to-one orthologs=0.58 and 0.59, coefficients of variation=0.29 and 0.30 respectively), marking syntenically stable regions. Together, these observations highlight chromosome-specific remodeling consistent with the structural divergence observed in our synteny analysis.

### Immune-related gene family from lineage-specific expansions

Building on the observed chromosome-scale heterogeneity in ortholog conservation and SV-associated remodeling, we systematically mapped lineage-specific expansion (LSE) segments across the *C. virginica* genome in direct comparison with *M. gigas*. Using joint filters on gene count and shared ortholog proportion (one-to-one and many-to-many), we identified 188 candidate 500 kb bins (17.7% of the genome), which later merged into 18 aggregated LSE segments (Figure 5A). These regions showed markedly reduced shared ortholog content with *M. gigas* (14.2%) (Figure 5B). Gene ontology (GO) analyses revealed significant functional enrichment in 8 of the 18 LSEs (in comparison to 8,553 overlapped and annotated genes in the genome background) (Data S6), with 24 immune-related terms enriched across five segments and elevated immune signals detected in 10 out of the remaining 13 segments (Figure 5C). These results highlight a genome-wide trend in *C. virginica* toward immune-related functional enrichment within LSEs in comparison with *M.gigas*.

**Figure 5.**
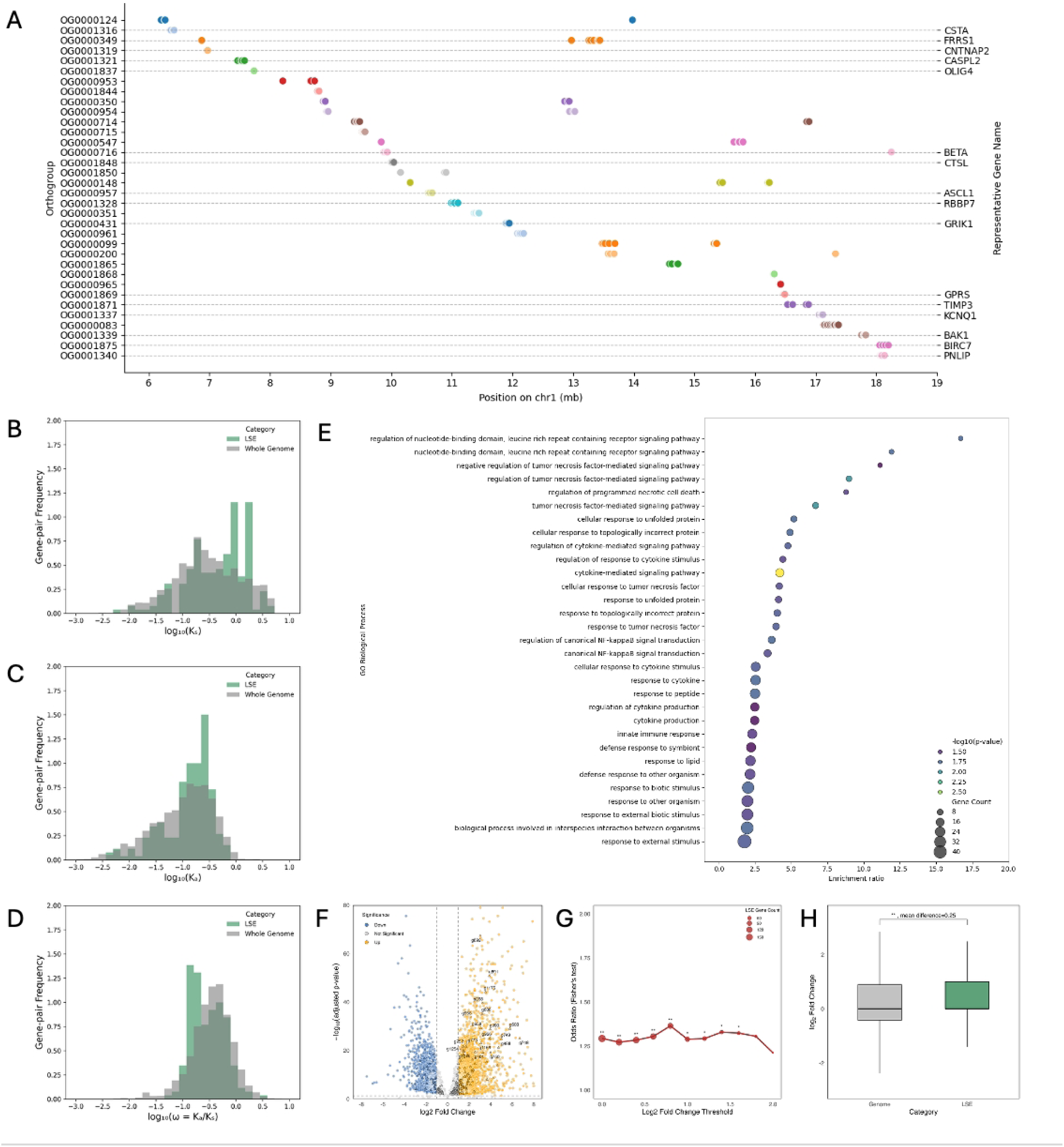
Genomic organization, functional enrichment, and evolutionary dynamics of the 13 Mb lineage-specific expansion (LSE) on *C. virginica* chromosome 1. **(A)**Spatial distribution of local gene families within the LSE, including only orthogroups with ≥3 C. virginica expanded members. Each dot represents a gene, colored by orthogroup, revealing a clustered organization of lineage-specific expansions. Gene families with known functional annotation are labeled on the right side. **(B)**Distribution of synonymous substitution rates (Ks) for paralogous gene pairs within the LSE. The prominent low-Ks peak indicates a recent burst of duplications. **(C)**Distribution of nonsynonymous substitution rates (Ka) for LSE paralogs compared to the genome-wide background, showing slightly elevated values in the LSE, consistent with lineage-specific diversification. **(D)**Distribution of nonsynonymous/synonymous substitution ratios (ω=Ka/Ks) for LSE genes versus genome-wide genes. LSE genes show a right-shifted distribution, consistent with relaxed constraint or lineage-specific adaptation. **(E)**Gene Ontology (GO) enrichment analysis of annotated genes within the LSE relative to the genome-wide background. Bars represent-log10 adjusted p-values, highlighting enrichment in immune, stress response, and signal transductin categories. **(F)**Volcano plot of differential gene expression under 2′3′-cUA treatment versus control. Circles represent genome-wide genes, and triangles highlight LSE genes, showing a strong overrepresentation of LSE genes among upregulated genes. **(G)**Line plot of odds ratio and significance for LSE gene enrichment among upregulated genes across increasing log2 fold-change thresholds. **(H)**Boxplot of log2 fold-change distributions comparing LSE genes to the genome-wide background. LSE genes exhibit significantly higher induction under immune stimulation.

To illustrate this trend in more details, we highlight the largest LSE detected: a 13Mb region on chromosome 1 (5,584,859-18,885,845) encompassing 908 genes. This interval coincides with a complex SV-associated gap, mostly entirely lacking one-to-one orthologs and dominated by *C. virginica*-specific and expanded families (Figure S1). Gene content was strongly enriched for lineage-specific (fold enrichment=2.60, *p*-value=8.90×10^-40^) unassigned (fold enrichment =2.72, *p*-value=3.96×10^-36^), and expanded orthogroups (fold enrichment =1.67, *p*-value=5.39×10^-10^), while one-to-one orthologs were nearly absent (fold enrichment =0.03, *p*-value=1.00). These features confirm the region as a concentrated hotspot of lineage-specific innovation.

Within the 13Mb lineage-specific expansion (LSE) of *C. virginica*, we identified 36 multi-copy gene families (≥4 members). To quantify their genomic organization, we calculated the coefficient of variation of pairwise gene distances and applied K-means clustering (k=2) to classify families as clustered (coefficient of variation≤0.92) or dispersed (coefficient of variation≥1.40) (Table S9). This analysis revealed 23 clustered families forming compact tandem arrays with high strand consistency, and 13 dispersed families spanning up to several megabases with mixed strand orientation. These contrasting architectures indicate that the region combines local tandem duplications with more complex multi-locus rearrangements, marking it as a structural hotspot shaped by extensive post-divergence remodeling relative to *M. gigas* (Figure 5D).

Analysis of amino acid substitution patterns within the 13Mb LSE revealed distinct evolutionary signatures compared to the genomic background (Figure 5E). Synonymous substitution rates (K_s_) were significantly higher in the LSE (Cliff’s δ=0.186, *p*-value=1.25×10^-5^) indicating that most duplications in this region originated after the divergence from *M. gigas* and represent relatively ancient events. Nonsynonymous substitution rates (K_a_) were also modestly elevated (Cliff’s δ=0.143, *p*=8.07×10^-4^), suggesting limited functional diversification among a subset of duplicates. In contrast, the overall ω (K_a_/K_s_) was slightly lower than the genomic background (Cliff’s δ=-0.137, *p*-value=1.26×10^-3^), consistent with pervasive purifying selection maintaining functional constraint across most copies. Collectively, these results indicate that the 13Mb LSE arose from ancient duplications, followed by modest functional diversification, with most retained duplicates subsequently constrained by purifying selection.

Among the 268 functionally annotated genes within the 13Mb LSE, GO enrichment revealed a strong focus on immune and stress-response processes (Figure S2). We further analyzed differential gene expression under 2′3′-cUA treatment using previously published RNA-seq data^26^ (8 treated vs. 8 control samples; Data S7). Among the 10,326 differentially expressed (DE) genes identified genome-wide, 302 were located within the LSE--a modest but statistically non-significant enrichment (odds ratio=1.065, *p*-value=0.213). Directional analysis revealed that 87 LSE genes were upregulated (log_2_(Fold Change)>1) and 33 were downregulated (log_2_(Fold Change) <1), compared to 2,462 upregulated and 1,624 downregulated genes genome-wide (Figure 5F). Notably, across a range of increasingly stringent log_2_(Fold Change) thresholds, LSE genes consistently showed significant enrichment of upregulated genes relative to the genomic background (Figure 5G), suggesting a coordinated transcriptional response. Furthermore, the permutation test confirmed a significantly elevated expression shift within the LSE (mean log_2_(Fold Change) difference=0.250; *p*-value=8.0×10^-4^) (Figure 5H), supporting the hypothesis that this lineage-specific expansion harbors stress-responsive genes under regulatory activation during immune challenge.

We examined gene family expansion events within this lineage-specific expansion (LSE). Among all functionally annotated families, the orthogroup containing FRRS1 showed the highest regional duplication count (n=8), with five members annotated as FRRS1. Four paralogs (g949, g950, g952, g954) form a compact tandem-like cluster (chromosome 1:13,283,141-13,446,823), whereas g488 (chromosome 1:6,871,498-6,877,572) resides near the LSE boundary. The FRRS1 cluster exhibits a spatial and evolutionary hierarchy (Figure 6A). Based on amino acid substitution estimates, g488, located at the LSE boundary, represents the most ancient copy (average K_s_=62.3, ω=0.01) (Figure 6B). The first interior duplicate, g954, diverged from g488 (K_s_ to g488=64.6), and subsequently gave rise to g952 (K_s_=5.93, ω = 0.11). The youngest tandem pair, g949 and g950 (pairwise K_s_=0.09, ω=0.36), arose from the most recent duplication within a topologically associated domain (TAD). This organization--ancestral duplication at the LSE boundary, interior expansion, and a recent tandem pair within the TAD--is reflected by decreasing K_s_ and increasing ω values, consistent with the multiple sequence alignment (MSA) based phylogenetic tree (Figure 6C). At the protein domain level, FRRS1 paralogs show progressive domain loss and rearrangement (Table S10). The ancestral copy, g488, retains the canonical Reeler-DOMON-Cytochrome b561 configuration (Figure 6D). The earliest interior duplicate, g954, gained an additional DOMON domain. The next duplicate, g952, lost the Reeler domain, marking the start of domain simplification. The most recent tandem duplicates continue this trajectory: g949 lacks the Reeler domain, while g950 has further lost a DOMON domain, retaining only Cytochrome b561. In aggregate, the FRRS1 paralogs exemplify how lineage-specific expansion can generate exceptional structural and domain diversity within a single genomic region of *C. virginica*.

**Figure 6.**
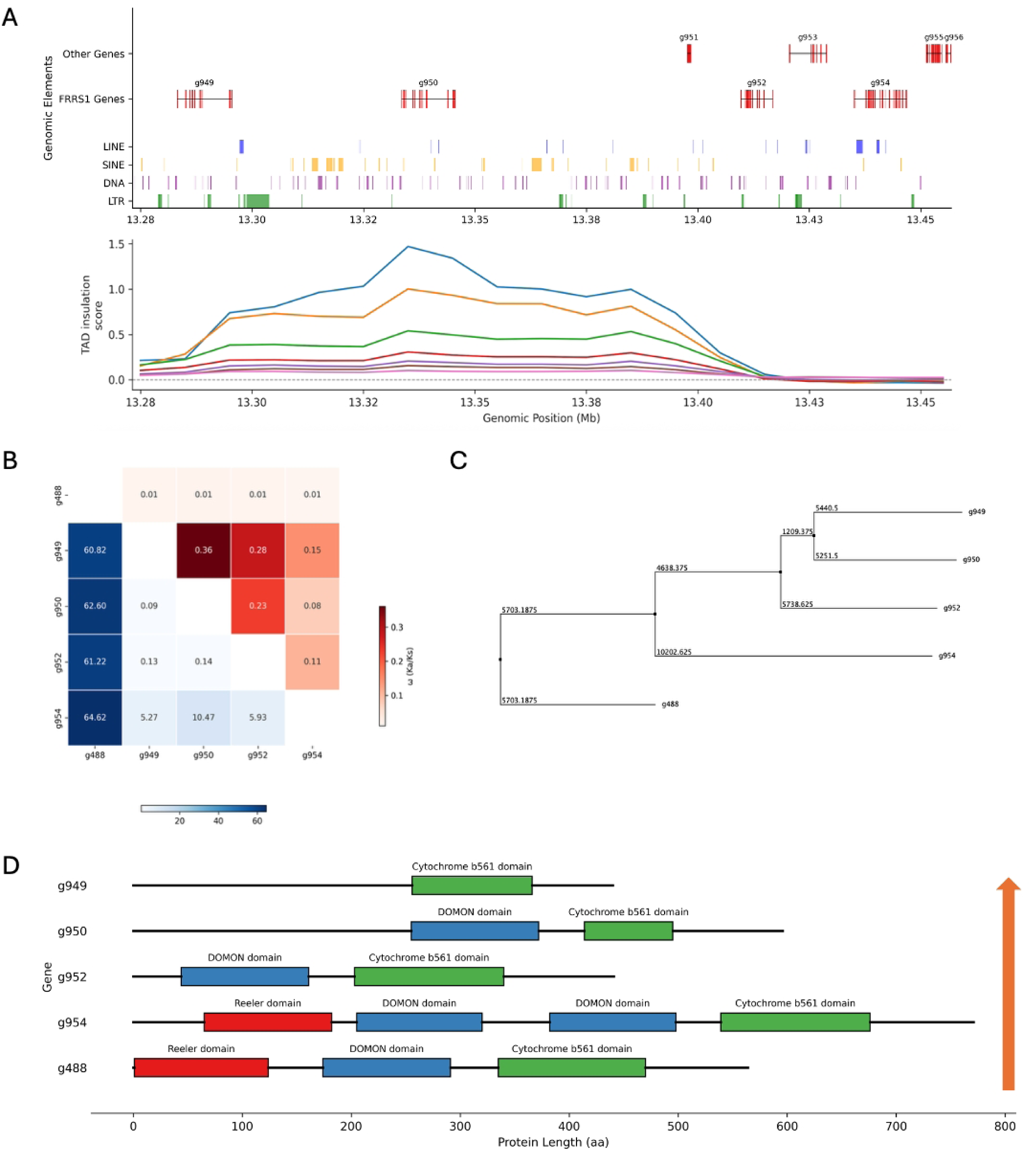
Local expansion and functional divergence of the FRRS1 gene family within the 13 Mb LSE. **(A)**Genomic landscape of the FRRS1 gene cluster within the LSE. The top major track shows local gene models, splitted into FRRS1 genes (lower level) and other genes (upper level). The second major track displays the distribution of transposable elements (LINE, SINE, DNA transposon, and LTR), while the bottom panel shows TAD insulation scores based on 10kb bins. **(B)**Pairwise evolutionary metrics for FRRS1 paralogs. The upper triangle shows ω (Ka/Ks) values, indicating the strength of selective pressure, while the lower triangle shows synonymous substitution rates (Ks), reflecting the relative timing of duplication events. **(C)**Neighbor-joining phylogeny of FRRS1 paralogs in the LSE, showing sequential duplication events within this gene family. **(D)**Protein domain architecture of FRRS1 paralogs in the LSE. Conserved domains include Reeler (red), DOMON (blue), and Cytochrome b561 (green). Several paralogs exhibit domain-level variations, suggesting potential subfunctionalization or pseudogenization following duplication.

## Discussion

In this study, we present an updated *C. virginica* genome assembly that markedly improves structural accuracy and completeness over the widely used reference (*C_virginica_3.0*), featuring extensive haplotig purging and chromosome misassemblies correction. By integrating Hi-C scaffolding, manual curation, and haplotig removal, we reduced duplicated BUSCOs from 18.7% to 1.1% while achieving 99.7% gene completeness, ensuring that genomic features such as structural variation and the 13Mb lineage-specific expansion reflect genuine biology rather than assembly artifacts. These improvements provide a robust foundation for downstream analyses, including comparative genomics, population studies, and functional investigations.

Comparative analyses across four oyster species confirm strict one-to-one chromosome correspondence, reinforcing long-recognized stability of chromosome content within *Ostreidae*, while resolving earlier reports of disrupted synteny^22,23^ that stemmed from assembly artifacts. Structural divergence among oysters is therefore driven primarily by intrachromosomal events--local shuffling, inversions, and lineage-specific insertions or deletions--rather than interchromosomal rearrangements. The extent of intrachromosomal remodeling, however, is highly uneven across chromosomes. For example, *C. virginica* chromosome 10 shows highly fragmented correspondence with extensive block shuffling, whereas chromosome 8 remains largely syntenic across species. These differences parallel disparities in repeat content and suggest that repeat accumulation may predispose certain chromosomes to structural instability, promoting recurrent reshuffling and the erosion of long-range collinearity, consistent with observations of repeat-driven structural dynamism in oysters^21,27–29^.

Lineage-specific expansions (LSEs) are a recurring feature of the *C. virginica* genome, many of which are enriched for immune-and stress-related pathways. Across the genome we identified 18 aggregated LSE segments, nearly half of which showed certain level of functional enrichment for immune processes, suggesting that oyster LSEs contribute disproportionately to diversification of defense and resilience mechanisms. This trend is consistent with previous reports of expanded immune receptors and stress-response genes in oysters^15,27,30^. Extending these observations, we uncovered a previously unreported immune megacluster--a striking 13Mb LSE on chromosome 1, flanked by a 4.4Mb colinear synteny block and a 3.3Mb inverted block relative to *M. gigas*. Such a configuration aligns with theoretical and empirical observations in which large inversions act as genomic islands, suppressing recombination and thereby enabling coordinated gene expansion^31,32^. The inversion on chromosome 1 may have provided a recombination-sheltered environment that promoted the retention and clustering of lineage-specific and expanded families, ultimately giving rise to an unusually contiguous block of gene innovation in *C. virginica*.

The structural organization of the 13Mb LSE, with tightly packed tandem clusters interspersed with dispersed paralogs, reflects a mixed history of local duplication and rearrangement. Inconsistent strand orientations further suggest that expansion was accompanied by short-range inversions and reshuffling. This dynamic architecture mirrors well-established models of immune gene evolution, such as the major histocompatibility complex (MHC)^33^ and immunoglobulin loci in vertebrates^34^. A representative example is the FRRS1 family, which has undergone extensive local duplication and domain remodeling. Proteins containing DOMON and Cytochrome b561 domains have been shown in other systems to regulate redox balance and modulate stress responses and iron homeostasis^35^, and the progressive erosion and reshuffling of these domains in FRRS1 suggest emerging functional divergence among closely linked paralogs. Similar domain-level remodeling has been described in plants, where Cytochrome b561-DOMON proteins acquired redox-regulatory roles through specialization^36^. Beyond FRRS1, the high density of unassigned or rapidly diverging genes within the 13Mb LSE may represent both a reservoir of oyster-specific innovations and remnants of decayed paralogs, consistent with the birth-and-death model of immune gene family evolution^33^.

Functionally, the enrichment of immune-and stress-related genes within this 13 Mb LSE, together with its strong transcriptional activation under 2′3′-cUA challenge, suggests that it serves as a flexible genomic reservoir for environmental adaptation. Responding genes in this region show significantly greater upregulation than the genomic background, consistent with potential co-regulation mechanisms that enable rapid, localized immune responses^27^. Such concentrated, inducible clusters are reminiscent of plant resistance gene superclusters, yet the sheer size and contiguity of this oyster LSE is unusual among animals, where immune loci are typically more fragmented^37,38^. This distinctive organization likely reflects the unique selective pressures faced by *C. virginica*, where high pathogen load and environmental variability favor the consolidation of specialized immune functions within a single genomic hub.

While this study provides a high-resolution view of genome structure and gene family evolution in *C. virginica*, several limitations should be noted. First, although the assembly substantially improves contiguity and accuracy relative to previous references, it still contains unresolved gaps in repeat-rich regions that may harbor functionally relevant elements. Our haplotig-purging strategy, while effective in reducing redundancy, may also have removed genuine paralogs, leading to conservative estimates of recent duplications. Second, because the assembly derives from a single individual, it may not capture the full spectrum of structural and gene content variation present across natural populations. Third, the 13 Mb LSE showed clear transcriptional activation under 2′3′-cUA stimulation, but not under other environmental stressors such as heat or *Perkinsus marinus* infection, based on reanalysis of available transcriptomes. This context-specific response suggests that its contribution may be stimulus-dependent, and its broader role in stress adaptation remains unresolved. Finally, while our analyses revealed strong associations between structural variation, gene family expansion, and immune-related activation, these relationships remain correlative. Functional validation through perturbation, regulatory assays, or long-read transcriptomics will be essential to establish causal links between genome structure and adaptive function.

In conclusion, our study shows that genome evolution in oysters is marked by long-term stability in chromosome content, coupled with uneven lineage-specific expansions that disproportionately involve immune and stress-response families. The 13Mb megacluster on chromosome 1 illustrates how such expansions can consolidate adaptive functions within a single genomic region, enabling coordinated transcriptional responses under immune challenge. These findings highlight lineage-specific expansion as a key mechanism of adaptive diversification in *Crassostrea virginica* and and provide a basis for comparative studies of genome evolution and adaptation in marine invertebrates. The reference genome presented here also provides the foundation for extending these insights to functional, ecological, and applied contexts.

## Methods

### Summary

Oysters were harvested from the Long Island Sound. After extraction DNA from the mantle tissues, we generated whole genome sequence data via Next Generation Sequencing, PacificBio, HiC technologies. We then assembled a reference genome. We identified single nucleotide polymorphisms (SNPs), as well as comparing to with the genomes of another species to assess known gene functions and inferring putative evolutionary history.

### Sample collection and Data Generation

All adult *Crassostrea virginica* individuals were collected from the same environment at a commercial oyster farm in Long Island Sound, New York. Immediately after harvest, specimens were placed on ice, transported to the laboratory, shucked, and whole tissues were flash-frozen in liquid nitrogen. Genomic DNA was extracted using the E.Z.N.A. Mollusca & Insect DNA Kit, and quality was assessed with NanoDrop spectrophotometry. Hi-C libraries were prepared with the Arima-HiC Kit according to the manufacturer’s protocol and sequenced on an Illumina NovaSeq X platform. Illumina whole-genome sequencing (WGS) libraries were prepared in parallel and sequenced on the same platform. High-molecular-weight DNA was also used to generate PacBio HiFi reads on the Revio system, following the standard PacBio protocol.

### Genome Assembly and Quality Control Assessment

We conducted genome assembly using raw data from PacBio HiFi long reads, Illumina whole-genome reads (WGS), and Arima Genomics Hi-C data. Raw Illumina reads were first assessed with FastQC v0.11.9 and adapter-and quality-trimmed using Trimmomatic v0.39 (Q30 cutoff) ^39^. PacBio HiFi reads were used directly without trimming. Primary contigs were assembled with Hifiasm v0.24.0 ^40^ using default parameters. To reduce allelic redundancy, one round of Purge_Dups v1.2.6 (default workflow) ^41^ was applied to the contig assembly, guided by Illumina and HiFi read depth profiles.

Chromosome-scale scaffolding was performed using Hi-C data. Trimmed Hi-C reads were aligned to contigs with BWA-MEM v0.7.17 ^42^, and scaffolding was conducted with Juicer v2.0.1 ^43^ and the 3D-DNA pipeline (2021-Mar-18 version) ^44^ under default parameters. Manual curation and misjoin correction were performed with Juicebox Assembly Tools v2.15. After scaffolding, residual haplotigs and unresolved heterozygous regions were identified using a two-step self-alignment approach. Whole-genome self-alignment was first performed with minimap2 v2.22 ^45^ and then confirmed with MUMmer4 v4.0.1^46^ (“nucmer --maxmatch-c 100-b 500-l 50”).

Candidate redundant regions were identified based on both alignment and coverage criteria. Non-redundant alignments were filtered to retain hits with alignment length ≥10kb and identity ≥95%. Illumina WGS coverage was calculated in 100kb bins, and half-coverage zones were defined as bins with coverage <65% of the genome-wide mean. Nearby half-coverage zones within 400kb were merged to delineate candidate heterozygous regions. A scaffold was considered for manual removal only if the residual haplotig segment was ≥500kb in length. Using this workflow, approximately 3.0Mb of residual haplotigs or unresolved heterozygous regions were manually removed, and scaffold breakends were stitched with 100 Ns at each join to represent unknown gap sequences, yielding the final non-redundant assembly. All remaining high-confidence self-alignments were classified into three categories based on their genomic positions: tandem duplications, where duplicated segments were separated by <100kb; segmental duplications, where duplicated segments were separated by >100kb on the same chromosome; and inter-chromosomal segmental duplications, where duplicated segments resided on different chromosomes.

Assembly quality was assessed using multiple complementary approaches. BUSCO v5.8.2 ^47^ with the mollusca_odb10 dataset evaluated gene completeness. Quast v5.3.0 ^48^ reported standard assembly statistics, including N50, L50, total size, and GC content. Merqury v1.4.1 ^49^ was used for 21-mer-based quality value (QV) estimation, with k-mer databases generated using Meryl v1.4.1 from both PacBio HiFi and Illumina WGS reads. Whole-genome structural consistency was further evaluated by comparing the new assembly to the previous *C. virginica_3.0* reference. Pairwise genome alignments were generated using MUMmer4 v4.0.1 (“nucmer --mum-c 100-b 500-l 50”), and alignment results were manually filtered to retain alignments ≥0.01Mb (10kb) and ≥ 90% identity. These filtered alignments were visualized as whole-genome dot plots, which revealed collinear chromosomes, scaffold-level misorientations, and large misjoins present in the previous assembly.

### Repetitive Element and Tandem Array Annotation

Repetitive elements in the C. virginica genome were identified entirely through de novo discovery. RepeatModeler v2.0.7 ^50^ was used to generate a species-specific repeat library. The assembly was then soft-masked with RepeatMasker v4.1.8 ^51^ using only the custom de novo library, classifying transposable elements (TEs) into major categories; including DNA transposons, LINEs, SINEs, and LTR retrotransposons. For quantitative analyses, overlapping TE annotations of the same class were merged into non-overlapping intervals to avoid double-counting bases covered by nested or partially overlapping repeats. Genome-wide TE content was calculated per chromosome and in 200kb non-overlapping windows, generating base-level coverage estimates for visualization and for correlation analyses with gene and duplication density.

Telomeric repeats were identified using Tandem Repeats Finder (TRF) v4.09.1^52^ with parameters “2 7 7 80 10 50 2000”. Candidate telomere tracts were defined as “TAAGGG” hexamer arrays of ≥60 bp, located within 10kb of chromosome ends, and showing ≥95% sequence identity. These telomeric sequences were generally moderate in length rather than forming long, uninterrupted blocks.

Candidate centromeric arrays were extracted from the TRF output using the following filters: total array length ≥10kb, repeat unit (period) ≥150 bp, copy number ≥10, and percent match ≥ 75%. The basic repeat units of the filtered arrays were then compared using BLASTN v2.13.0+^53^ (-evalue 1×10^-5^), and hits were retained if identity was ≥80% and aligned length ≥70% of the smaller array. Arrays passing these criteria were clustered into families by connecting arrays as nodes in an undirected graph when a qualifying BLASTN hit existed between them. These graph-based clusters were then manually reviewed to confirm family assignments and calculate centromere indices for each chromosome, which were compared against expected positions from prior C. virginica cytogenetic studies. Arrays forming family-specific clusters in chromosomal interior regions were annotated as putative centromeric repeats.

### Gene Prediction and Functional Annotation

Protein-coding genes were predicted using RNA-seq–guided annotation. Publicly available *C. virginica* RNA-seq datasets from 10 samples spanning mantle, gonad, and mixed tissues from both sexes were preprocessed with Fastp v0.20.0^54^ (“--cut_front --cut_tail --detect_adapter_for_pe --cut_mean_quality 20-l 50”). Filtered reads were aligned to the genome using STAR v2.7.11b^55^ in two-pass mode to improve splice junction detection. All alignments were then merged with Samtools v1.6^56^ to generate a unified BAM file for gene model training.

Gene structure prediction was performed using BRAKER v3.0.8^57^, which integrates RNA-seq alignments with ab initio modeling to generate high-confidence gene models. Coding sequences (CDS) and predicted proteins were extracted using GffRead v0.12.7^58^ to produce FASTA files for downstream analysis. Functional annotation was conducted with EggNOG-mapper v2.1.12^59^, which assigned orthology-based functional descriptions, putative gene names, Gene Ontology (GO) terms^60^, and KEGG pathways^61^.

For synteny analysis, predicted proteins (longest variant for each gene) were extracted using SeqKit 2.8.1^62^ and compared against the previous *C. virginica* reference genome using BLASTP v2.15.0+ (“-evalue 1e-5-max_target_seqs 5”). Collinear blocks were identified using MCScanX v1.0.0 ^63^ with parameters optimized for high-confidence synteny detection. A minimum of five consecutive anchor gene pairs was required to define a syntenic block, and gaps of up to 25 genes were allowed between adjacent anchors. Collinear gene pairs were scored with a base match score of 50, and a penalty of-1 was applied for each missing gene in a gap. An overlap window of five genes was used to merge nearby alignments, and gene pairs were required to pass a BLASTP E-value threshold of 1×10^-5^. Anchor gene pairs from MCScanX were subsequently used for genome-wide synteny visualization and for validating structural consistency between the new assembly and the previous reference.

Whole-genome sequencing reads from 19 additional *C. virginica* individuals were preprocessed and aligned to the new assembly following the same workflow as the reference sample. Aligned reads were later processed with SAMtools v1.12 to sort, remove PCR duplicates, and filter alignments with a mapping quality threshold of MQ≥30. We then followed the GATK v4.2.0.0 germline short variant discovery best practices pipeline^64^ to generate a SNP dataset from the 19 additional *C. virginica* individuals. Joint variant calling was performed across all samples, and raw SNPs were subjected to recommended hard filters to retain only high-confidence biallelic SNPs. Specifically, sites flagged by GATK as failing any of the following thresholds were removed: “QD<2.0”, “QUAL<30.0”, “SOR>3.0”, “FS>60.0”, “MQ<40.0”, “MQRankSum<-12.5”, or “ReadPosRankSum<-8.0”. Joint variant calling across the 20 resequenced individuals and the reference sample yielded 33,599,564 biallelic SNPs after GATK hard filtering.

To derive a conservative set of fixed differences between the reference individual and all others, we applied additional filtering steps. First, sites with missing genotype calls in the reference sample (n=159,592) or homozygous reference genotypes across all samples (n=2,927) were removed. We then excluded loci with a genotyping rate <0.8 across individuals. Finally, we identified positions where the reference sample was homozygous reference and all non-reference individuals were either homozygous alternate or missing. This strict criterion yielded a final set of 34,810 high-confidence SNPs.

### Comparative Synteny Analysis Among Oyster Species

The reference genome of *M. gigas* (xbMagGiga1.1), *M.angulata* (ASM2561291v2), *O. edulis* (xbOstEdul1.1) and their corresponding GFF annotation files were downloaded from NCBI. For each gene, the longest protein isoform was extracted to generate a non-redundant protein set. Pairwise BLASTP searches were performed between *C. virginica* and each oyster protein set using the same parameters as for intra-species synteny analysis, and collinear blocks were identified using MCScanX under the same settings.

Global gene family turnover was estimated using CAFE5 ^65^, which models gene family evolution under a stochastic birth-death process. The orthogroup count table from OrthoFinder v2.5.5 ^66^, and the species tree inferred from single-copy orthologs were used as input. A global lambda (birth/death rate) model was applied, and significantly expanded or contracted gene families were identified using a p-value threshold of 0.05. Expansion and contraction counts were summarized per species and visualized using stacked bar plots.

For in-depth comparative analysis between *M. gigas* and *C. virginica*, only synteny blocks containing ≥10 anchor genes were retained to ensure high-confidence structural correspondence. Potentially overlapping synteny pairs were resolved using a two-dimensional merging approach, where adjacent blocks overlapping in both species were merged to generate a non-overlapping final synteny set.

To ensure a consistent orientation for comparative analysis, *M. gigas* chromosomes were examined for potential strand inversions. Orientation correction was determined using a combined approach that considered both the cumulative length of forward versus reverse alignments and the sequential arrangement of synteny blocks along each chromosome. Chromosomes for which the majority of aligned sequence length and block ordering supported a reverse orientation were flipped in silico before downstream analysis. This procedure resulted in the orientation correction of chromosome 4,6,7,8, and 9 of *M.gigas*, improving both visual collinearity and the accuracy of quantitative synteny metrics.

To quantify large-scale chromosomal structure and rearrangements, synteny blocks were sorted by their order on the *C. virginica* genome and traversed sequentially. A flip count was incremented whenever the orientation of consecutive *M. gigas* blocks switched relative to *C. virginica*. Spearman rank correlations were then calculated between the ordered positions of *C. virginica* synteny block and their corresponding *M. gigas* positions to quantify global collinearity, with lower absolute correlations indicating more extensive chromosomal shuffling.

### Comparative Structural Variant Classification between *C. virginica* and *M. gigas*

Structural variants (SVs) between *C. virginica* and *M. gigas* were identified using anchor gene synteny derived from high-confidence MCScanX blocks (≥ 10 anchors), with *C. virginica* treated as the reference genome. Anchor genes were first sorted by their genomic positions on C. virginica, and the genome was traversed sequentially to analyze the arrangement of the corresponding *M. gigas* anchors. For each pair of consecutive *C. virginica* anchors, the inter-anchor distances were calculated as CV gap in *C. virginica* and MG gap in *M. gigas*. In total, 11,394 anchor genes produced 11,375 adjacent anchor pairs, and 24,542 non-anchor genes were also mapped for synteny context.

The distributions of CV gap and MG gap were examined to distinguish typical intergenic spacing from expanded intervals indicative of structural rearrangements. Large gaps were defined using the 99th percentile of the adjacent-anchor gap distributions, corresponding to 252,389 bp for CV gap and 285,688 bp for MG gap. These thresholds provided an empirical basis for SV classification. Intervals where both gaps were below the thresholds were considered trivial, representing normal intergenic spacing. When CV gap exceeded the threshold while Mg gap remained small, the interval was classified as a deletion in *M. gigas*. Local insertions in *M. gigas* were defined by a large MG gap and a small CV gap with no additional *C. virginica* syntenic genes present in the expanded Mg region, whereas foreign insertions in *M. gigas* were similarly defined but involved the presence of other *C. virginica* syntenic genes within the MG gap, indicating insertion of a segment from a different syntenic block. Inversions were identified when both gaps were small but the *M. gigas* anchors were in reversed orientation relative to C. virginica. All remaining cases, including intervals with large gaps in both genomes, mixed orientations, or overlapping rearrangements, were classified as complex SVs.

This gap-and orientation-based framework provided a mutually exclusive SV classification that distinguished local insertions, foreign insertions, deletions, inversions, and complex events, while consistently using *C. virginica* as the reference for comparative analysis.

### Orthogroup Classification and Structural Variation Colocalization Analysis between *C. virginica* and *M. gigas*

Orthologous gene groups were inferred using OrthoFinder between *C.virginica* and *M.gigas*, with the longest protein isoform selected for each gene in *C. virginica* and *M. gigas*. Default settings were applied for all-vs-all sequence similarity searches and orthogroup clustering. The resulting orthogroups were classified heuristically into seven categories for downstream analyses: one-to-one (single-copy genes in both species), many-to-many (multiple genes in both species), CV-specific (genes unique to *C. virginica*), MG-specific (genes unique to *M. gigas*), CV expansion (orthogroups with two or more *C. virginica* genes and a single *M. gigas* gene), MG expansion (orthogroups with two or more *M. gigas* genes and a single *C. virginica* gene), and unassigned (genes with no detectable orthology). Each *C. virginica* gene was annotated with its orthogroup category for integration with structural variant analysis.

To investigate the relationship between gene family evolution and structural variation, each *C. virginica* gene was mapped to the nearest SV interval, defined as the region between adjacent anchor genes classified as a deletion, local insertion, foreign insertion, inversion, or complex SV. For each SV interval, we quantified the number of genes in each orthogroup category and compared these distributions with genome-wide background intervals to assess enrichment or depletion of specific orthogroup types.

We also assessed transposable element (TE) content in relation to SVs. For each SV interval, TE content was measured within 50kb flanking the gap on both sides, providing the local repeat context for each structural variant. These values were compared against genome-wide background TE levels, calculated in non-overlapping 50kb windows, to determine whether SV regions are preferentially enriched for specific TE classes. This framework enabled the identification of lineage-specific gene family expansions and repeat-rich regions that colocalize with local structural rearrangements in *C. virginica*.

### Identification and functional characterization of *C.virginica* genome-wide LSEs

We characterized lineage-specific expansions (LSEs) across the *C. virginica* genome in direct comparison with *M. gigas*. First, the genome was partitioned into non-overlapping 500kb bins, and each bin was evaluated for (i) the total number of annotated genes and (ii) the proportion of genes with one-to-one or many-to-many orthologs shared with *M. gigas*. Bins with fewer than 10 genes were excluded. A depletion threshold for shared ortholog proportion was defined as one standard deviation below the genome-wide mean. Bins that passed both filters were classified as candidate LSE bins. Adjacent candidate bins were then merged into aggregated LSE segments, retaining only runs spanning at least three consecutive bins.

Functional enrichment of genome-wide LSEs was performed using annotated genes within each aggregated segment as the test set and all 8,553 annotated genes across the genome as the background. Gene Ontology (GO) enrichment analysis was carried out in the Biological Process ontology using a hypergeometric framework. Terms were tested against the Homo sapiens GO reference database, and p-values were corrected using the false discovery rate (FDR). Enrichment results were summarized across all 18 aggregated LSEs, with special emphasis on immune-related terms as defined by a curated immune GO universe. Specifically, a set of 12 anchor GO terms directly relevant to immune processes (GO:0002376, GO:0006952, GO:0006954, GO:0019221, GO:0034097, GO:0045087, GO:0002250, GO:0019882, GO:0002682, GO:0070098, GO:0050900, GO:0030595) was defined a priori. All descendant terms of these anchors were collected from the GO hierarchy. In total, 1,773 GO biological process terms were included in the immune panel. Significant enrichment of immune terms was then evaluated segment by segment, and the global distribution of enrichment across all LSEs was summarized.

### Orthogroup Mapping, Functional Analysis, and Evolutionary Dating of the 13Mb Lineage-Specific Expansion

The 13Mb lineage-specific expansion (LSE) in *C. virginica* was defined based on synteny breakpoints, spanning chr1:5,584,859-18,885,845, and delimited by the anchor genes g373 and g1280, which mark the boundaries between two adjacent synteny blocks in the *C. virginica*-*M. gigas* alignment. This interval represents a *C. virginica*-specific insertion with no colinear counterpart in *M. gigas*.

All *C. virginica* genes within the LSE were annotated with orthogroup categories inferred using OrthoFinder v2.5.5 and the longest isoform per gene: one-to-one, many-to-many, *C. virginica*-specific, *C. virginica*-expanded, *M. gigas*-expanded, and unassigned. By definition, *M. gigas*-specific orthogroups are absent in this region, and *M. gigas*-expanded orthogroups were extremely rare (n=6) and were excluded from downstream analysis. This orthogroup mapping formed the basis for assessing duplication architecture and functional organization in the LSE.

Gene family organization within the LSE was analyzed for 36 orthogroups containing at least four *C. virginica* genes. For each orthogroup, all pairwise genomic distances among its genes were used to calculate the mean distance, standard deviation, and coefficient of variation as metrics of spatial dispersion. A strand consistency score--the proportion of all gene pairs sharing the same transcriptional orientation--was computed to distinguish tandem-like clusters from irregularly oriented duplications. K-means clustering (k=2, random seed=42) applied to standardized mean distance and coefficient of variation values classified orthogroups into compact, tandem-like clusters or dispersed duplications, highlighting the heterogeneity of duplication architecture within the LSE.

For the focal 13Mb LSE on chromosome 1, functional interpretation was performed at the orthogroup level. Orthogroups were assigned functional annotations based on the most frequent EggNOG-mapper annotation among their member genes. Enrichment analysis used the set of 268 unique LSE gene names as the foreground and the same genome-wide background of 8,553 annotated genes. As with the genome-wide analysis, enrichment was performed in the Biological Process ontology using Fisher’s exact test and FDR correction.

To estimate the evolutionary timing of gene duplications, we performed synonymous substitution (Ks) analysis using wgd v2 ^67^. Coding sequences (CDS) of all LSE genes were compared to genome-wide paralogs to generate Ks distributions. To ensure robust estimates, we filtered pairwise alignments to remove low-quality matches, requiring dN (nonsynonymous substitutions per site), dS (synonymous substitutions per site), and dN/dS values between 0 and 10, an effective aligned length (alignment length×coverage) of at least 150 bp, and an alignment identity of at least 30%. Pairs passing these filters were used to construct the LSE-specific Ks distribution and to compare it with the genome-wide background, enabling us to distinguish recent from ancient duplication events.

### RNA-seq Processing and Differential Expression Analysis

Previously published RNA-seq data from Crassostrea virginica exposed to 2′3′-cUA treatment were downloaded from the NCBI Sequence Read Archive (Project ID:*PRJNA970795*). Raw paired-end reads were quality-controlled using fastp v0.23.2, with removal of Illumina adapters, polyG tails, and low-quality bases (Q30 cutoff). Clean reads were aligned to the newly assembled C. virginica reference genome using STAR v2.7.11a under default settings. Gene-level read counts were generated using featureCounts from Subread v2.0.3 ^68^ based on the custom GFF3 annotation file.

Downstream analysis was performed in R v4.4.2 with DESeq2 v1.46.0 ^69^. Genes with normalized read count smaller than 10 were filtered out, and variance-stabilizing transformation (VST) was applied for visualization and principal component analysis. Differential expression between treated and control samples was assessed using Wald tests with Benjamini-Hochberg multiple-testing correction. Genes were considered differentially expressed if they had an adjusted p-value <0.05 and |log₂ fold change|>1 (log_2_FC>1 for upregulation, log_2_FC<-1 for downregulation). For visualization, extremely low-effect genes were filtered using |log_2_FC|>1×10^-4^ to remove numerical noise.

To evaluate whether the 13Mb LSE on chromosome 1 shows distinct transcriptional responses, we performed permutation testing with 10,000 random samplings (random seed=42) comparing the mean log_2_FC of LSE genes versus genome background genes (defined as all genes outside the 13Mb LSE). This allowed us to assess whether expression changes in the LSE significantly deviated from genome-wide expectations.

### Evolutionary History Reconstruction of the FRRS1 Gene Family

The FRRS1 gene family, which exhibits pronounced expansion within the 13Mb LSE, was analyzed to reconstruct its duplication history, structural context, and expansion mechanism. All FRRS1 orthogroup members were mapped to the final *C. virginica* assembly, and their gene models, including intron/exon structure, were defined using the longest isoform per gene. These metrics distinguished tandem-like, transposed, and dispersed duplications, revealing a stepwise accumulation pattern with older duplications near LSE boundaries, intermediate transposed copies, and the youngest tightly clustered tandem duplicates.

Duplication timing and selective pressures were assessed first using pairwise Ks and Ka/Ks analysis, following the same workflow and filtering criteria as the genome-wide study. These analyses differentiated ancient duplications from recent expansions and evaluated whether FRRS1 members are evolving under purifying or relaxed selection, providing a temporal framework for subsequent phylogenetic interpretation.

To examine the structural context of expansion, local insulation scores were calculated using HiCExplorer v3.7.2 ^70^ on 25kb genomic bins, with Hi-C reads remapped to the final assembly. Scores were computed with step=50kb, minDepth=100kb, maxDepth=500kb, and binsize=25kb, quantifying local chromatin contact insulation and identifying potential topologically associating domain (TAD) boundaries. This analysis tested whether the youngest FRRS1 duplications are positioned in structurally insulated regions that promote stable tandem duplication.

For evolutionary reconstruction, coding sequences of the longest isoforms were aligned using MAFFT v7.525 ^71^, and a profile hidden Markov model was generated with HMMER v3.4 ^72^ to refine homologous positions. A distance-based neighbor-joining tree was then built in Jalview v2.11.4.0 ^73^ to visualize duplication chronology. Protein functional domains were annotated using InterPro 106.0 ^74^ across all available databases, with overlapping hits resolved by prioritizing the annotation with the lowest E-value, ensuring confident functional assignments based on the longest protein isoform of each gene.

## Statistical Analysis

All statistical analyses were performed in Python (v3.12) using the SciPy and statsmodels packages, and RNA-seq differential expression analyses were conducted in R (v4.4.2) with DESeq2 (v1.46.0). Two-sided Mann-Whitney U tests were used for non-parametric comparisons, including gap sizes, gene counts, repeat densities, orthogroup distributions, and pairwise Ka, Ks, and Ka/Ks (ω) distributions. Fisher’s exact tests were applied for categorical enrichment analyses--including Gene Ontology (GO) enrichment and orthogroup overrepresentation near structural variation breakpoints--and for testing differential expression (DE) enrichment of LSE genes relative to the genome background. Spearman correlation was used to assess synteny block ordering concordance between *C. virginica* and *M. gigas*, while Pearson correlation was used to evaluate the relationship between chromosomal gene density and repeat content. For RNA-seq analyses, LSE versus background expression shifts were evaluated using a 10,000-iteration permutation test (seed=42). Benjamini-Hochberg false discovery rate (FDR) correction was applied for GO enrichment and DE analyses to account for multiple comparisons. Unless otherwise specified, statistical significance was defined as p < 0.05.

## Declarations

We confirm that all experiments were approved by the IACUC committee at Yale University. We confirm that all experiments were performed in accordance with IACUC guidelines and regulations at Yale University. We ensure that our manuscript reporting adhered to the ARRIVE guidelines for the reporting experiments. We confirm that our methodology can be fully evaluated and utilized.

## Ethics approval and consent to participate

-This work is exempt from the IACUC protocol.

-The name(s) of the Institutional Review Board(s) or Ethics Committee(s): Professor Peter Smith, DVM at Yale University.

## Consent for publication

Not applicable.

## Availability of data and materials

All data are contained in the manuscript, the supplementary file (will be submitted to SRA.) Accession number of the reference whole genome sequence data will be available. The raw whole genome sequencing data generated by the Hoh lab will be available on the public domain.

## Authors’ contributions

Conceptualization: JH

Methodology: ZL, ZC, IT, BK, AD, JH

Investigation: ZL, ZC, JH

Visualization: ZL, JH

Funding acquisition: JH

Project administration: ZL, ZC, IT, JH

Supervision: ZC, IT, BK, AD, JH

Writing - original draft: ZL, JH

Writing - review & editing: ZL, LC, AP, WB, ZC, AD, JH

## Acknowledgements

We thank the team at the Oyster Farming Technologies.

